# Disentangling the effects of jasmonate and tissue loss on the sex allocation of an annual plant

**DOI:** 10.1101/2021.11.10.468055

**Authors:** Nora Villamil, Benoit Sommervogel, John R. Pannell

## Abstract

Selection through pollinators plays a major role in the evolution of reproductive traits. However, herbivory can also induce changes in plant sexual expression and sexual systems, potentially influencing conditions governing transitions between sexual systems. Previous work has shown that herbivory has a strong effect on sex allocation in the wind-pollinated annual plant *Mercurialis annua*, likely mediated by resource loss. It is also known that many plants respond to herbivory by inducing signalling, and endogenous responses to it, via the plant hormone jasmonate. Here, we attempt to uncouple the effects of herbivory on sex allocation in *M. annua* through resource limitation (tissue loss) versus plant responses to jasmonate hormone signalling. We used a two-factorial experiment with four treatment combinations: control, herbivory (25% chronic tissue loss), jasmonate, and combined herbivory and jasmonate. We estimated the effects of tissue loss and defence-inducing hormones on reproductive allocation, male reproductive effort, and sex allocation. Tissue loss caused plants to reduce their male reproductive effort, resulting in changes in combined sex allocation. However, application of jasmonate after herbivory reversed its effect on male investment. Our results show that herbivory has consequences on plant sex expression and sex allocation, and that defence-related hormones such as jasmonate can buffer the impacts. We discuss the physiological mechanisms that might underpin the effects of herbivory on sex allocation, and their potential implications for the evolution of plant sexual systems.

## Introduction

Flowering plants are remarkable for the diversity of their reproductive structures (flowers and inflorescences) and their sexual systems, which range from simultaneous hermaphroditism to completely separate sexes (dioecy), including intermediate systems in which males or females co-exist with hermaphrodites (Barrett, 2002). Much of this diversity is due to genetic differences that have accumulated between populations and species, but plant sexual systems are also often modulated by plastic responses of individuals’ sex expression to environmental or status-related cues, i.e., they reflect norms of reaction to their environment (Barrett, 2002, Cossard and Pannell, 2019, López and Domínguez, 2003). Pollinator-related selection is thought to be a major agent shaping both fixed and plastic aspects of plant sexual systems, but the influence of antagonists has probably been underestimated (Carr and Eubanks, 2014, Lucas-Barbosa, 2016, Santangelo et al., 2019, Johnson et al., 2015, Strauss et al., 1996). Herbivory, for instance, can affect floral display (Strauss et al., 1996, Santangelo et al., 2019), colouration (Strauss et al., 2004, Vaidya et al., 2018), floral scents (Ramos and Schiestl, 2019, Kessler et al., 2011), sex ratios (Hendrix and Trapp, 1981, Krupnick and Weis, 1998, Krupnick et al., 2000, Thomson et al., 2004) and plant sexual expression (Villamil et al., 2021). Moreover, selection on plant mating systems (e.g., through expression of self-compatibility/self-incompatibility) can influence the evolution of anti-herbivore defences (Campbell and Kessler, 2013). Recent studies have exposed links between herbivory and aspects of plant mating, yet how this interaction affects the sex allocation in general and the expression of hermaphroditism versus dioecy specifically remains poorly understood.

Herbivory has at least two consequences for plants. Most directly, it causes tissue loss, reducing plant size and limiting resources available for allocation to maintenance, growth and reproduction through male and female sexual functions (Ivey and Carr, 2005, Mothershead and Marquis, 2000, Karban and Strauss, 1993, Lehtilä and Strauss, 1997, Lehtilä and Strauss, 1999, Poveda et al., 2003, Hambäck, 2001, Kessler et al., 2011). If herbivory influences the fitness gained through male versus female allocation differently, then selection should favour a norm of reaction that we might label ‘conditional sex allocation’ (Wang et al., 2021, Hirata et al., 2019, Freeman et al., 1980, López and Domínguez, 2003). Conditional sex allocation can imply complete sex changes when the fitness gains from the male or female sexual functions change with age or size (Ghiselin, 1969, Charnov, 1982), with selection favouring individuals that reproduce first through the sex whose reproductive value increases more slowly with size, and that then change to the other sex when they reach the age/size point at which fitness gains increase more rapidly (West, 2009). Norms of reaction that entail complete switches in gender from male to female are known from some plants, e.g., jack-in-the-pulpit (Bierzychudek, 1984b, Bierzychudek, 1984a, Policansky, 1987), in which individuals respond to their size (and presumably their resource status), but changes in sex allocation in modular organisms like plants is often more quantitative, with a more continuous shift in sex allocation from an emphasis on one function to an emphasis on the other (de Jong and Klinkhamer, 1989, de Jong et al., 1992, Klinkhamer et al., 1997, de Jong and Klinkhamer, 2005, Pannell, 1997b, Pannell et al., 2008). The reductions in plant size brought about by herbivory are thus more likely to elicit quantitative norms of reaction in sex allocation.

Herbivory also affects plants indirectly by triggering responses that help to prevent further attacks and to restore physiological equilibrium after damage. These responses can be endogenous, via electric, osmotic or hormonal signals within the plant (Farmer et al., 2020, Salvador-Recatalà et al., 2014), or exogenous, triggered by volatile signals from neighbouring leaves/plants (e.g., jasmonate) (Ballaré, 2011, Heil and Bueno, 2007b, Heil, 2009) or animals (e.g., herbivore eggs or saliva) (Anastasaki et al., 2015, Hilker and Meiners, 2002, Mumm et al., 2003), which are perceived through leaf stomata. Signalling mediated by the volatile hormone jasmonate is particularly interesting in the present context because it is known to play a key role in both defence responses and the regulation of sex expression. On the one hand, jasmonate is released by damaged plant tissues and thus functions as an air-borne signal of nearby herbivore activity that induces anti-herbivore responses in leaves that perceive it (Ballaré, 2011, Thaler et al., 2001, Heil and Bueno, 2007b, Heil and Karban, 2010); such a response may be elicited in yet undamaged leaves of the same plant, but neighbouring plants can also ‘eavesdrop’ on jasmonate signalling and induce their defences in preparation for likely future damage (Heil and Bueno, 2007a, Heil and Bueno, 2007b, Karban, 2008, Karban et al., 2012). On the other hand, jasmonate is known to play a role in regulating sex expression, flower development, and sexual differentiation (Wasternack et al., 2013, Yuan and Zhang, 2015, Cai et al., 2014, Acosta et al., 2009, Yan et al., 2012) as part of a hormonal crosstalk network interacting with cytokines, auxins and other hormones (Yuan and Zhang, 2015, Naseem et al., 2015, Wasternack et al., 2013, Robert-Seilaniantz et al., 2011). However, jasmonate effects on the female or male components of sexual development seem to be species-specific (Wasternack et al., 2013): in *Arabidopsis thaliana* (Browse, 2009) and in maize (Yan et al., 2012), jasmonate is essential for male flower development; in tomato it is required for female flower development (Li et al., 2004); and in rice jasmonate determines the sex of the developing sexual organs (Acosta et al., 2009, Yuan and Zhang, 2015, Cai et al., 2014).

In this study, we explored norms of reaction in sex expression and sex allocation to herbivory in an experiment designed to uncouple its direct (through tissue loss) and indirect effects (due to defensive jasmonate signalling). Classic plant resource allocation theory suggests that increased investment in reproduction and growth comes at a cost in allocation to defence, and *vice versa* (Herms and Mattson, 1992). We tested the hypothesis that herbivory should cause plants to reduce their reproductive effort through a trade-off between resources allocated to defence versus reproduction. We reasoned that defence-related traits, such as those induced by jasmonate signalling, may link the coordinated evolution of reproductive and defensive traits. Growing evidence suggests that reproductive and defensive traits are not independent (Campbell, 2015, Carr and Eubanks, 2014, Lucas-Barbosa, 2016, Johnson et al., 2015), yet the effects of plant defensive strategies on trade-offs between male and female plant fitness and sex allocation still remain largely unexplored (Garcia and Eubanks, 2019).

To uncouple the direct and indirect effects of herbivory on the sex expression and to test the role of jasmonate on conditional sex allocation, we conducted a two-factorial experiment manipulating tissue loss (25% chronic defoliation) and plant anti-herbivore defences via the jasmonate pathway (external application of jasmonate), and measured sexual expression in plants with both a male and a female function. In many species (Reymond and Farmer, 1998, Wang et al., 2019, Han, 2016), plants that are damaged automatically release jasmonate into the environment, and it is therefore impossible for them to experience tissue loss without jasmonate release. However, our experimental design included all other biologically possible combinations: un-manipulated control plants, plants subject to tissue loss only, plants subject to jasmonate only, and plants subject to tissue loss and jasmonate together.

Our experiment used females of the wind-pollinated annual dioecious plant *Mercurialis annua* that have evolved greatly enhanced leaky sex expression following the experimental removal of all males from their populations, as a result of selection for reproductive assurance and greater siring success (Cossard and Pannell, 2021, Cossard et al., 2021). Leaky sex expression (the production of gametes of the opposite sex by plants with separate sexes) is common in dioecious plants and likely plays an important role in reversions from dioecy to hermaphroditism (Cossard and Pannell, 2019, Ehlers and Bataillon, 2007). Because the genotypes used in our experiment are females that are now expressing a greatly enhanced leaky male flower production that has evolved in the absence of herbivory, we had no *a priori* expectation for the how they should respond to simulated herbivory. However, previous work on dioecious *M. annua* has shown that simulated herbivory enhances leaky sex expression in both males and females (Villamil et al., 2021), for reasons that may or may not be adaptive. We might thus have expected this tendency for greater leakiness under herbivory to have been retained or even enhanced in the genotypes we used in our experiment. Surprisingly, we found that females with leakiness enhanced through experimental evolution actually reduced their male flower production in response to tissue loss in our experiment. Equally surprising, this response to simulated herbivory was erased in plants that were exposed to hormone signalling. Our experiment thus reveals complex norms of reaction to herbivory that, by definition, cannot be adaptive in their detail, but that represent intrinsic pleiotropic responses to prior selection on reproductive allocation.

## Materials and methods

### Study system

*Mercurialis annua* (Euphorbiaceae) is an annual wind-pollinated herb distributed throughout central and western Europe and around the Mediterranean Basin (Tutin et al., 1968). The species has long been used as a model system to investigate the evolution of dioecy and sex expression (Yampolsky, 1919, Yampolsky, 1930, Obbard et al., 2006, Pannell, 1997b) due to its great diversity and plasticity in sex expression and sexual systems (Yampolsky, 1930, Pannell, 1997a, Cossard et al., 2021, Cossard and Pannell, 2021). *M. annua* has chromosomal sex determination (XX♀; XY♂) (Russell and Pannell, 2015), mediated by endogenous hormonal signalling. Males are feminised by exogenous application of cytokinins, and females are masculinised by auxins (Durand and Durand, 1991). The plants used in this study are monoecious XX females that have recently evolved pollen production under natural selection under experimental evolution (Cossard et al., 2021) and are therefore now functionally hermaphroditic. These plants are an ideal system in which to test the effects of herbivory on sex allocation for two reasons. First, although *M. annua* has chromosomal sex determination, its sex expression is also hormonally regulated. Second, these plants have highly plastic sex allocation, which has evolved very recently; therefore, we expect the interference of defensive signals on sex allocation, if it occurs, to reflect deeply conserved physiological reaction norms that have not been shaped by recent natural selection.

### Plant culturing

Seeds generated through a selection experiment (Cossard et al., 2021) were sown in plastic trays with sterilised soil (Ricoter 163 soil) and kept under stable greenhouse conditions (25°C, 50% humidity; October 2019) at all times. When seedlings had flushed their first four leaves, approximately three weeks after sowing, they were transplanted into individual pots (Teku serie TO 14 D) with soil (Ricoter 140 soil) and slow-release fertilizer (Hauert Tardit: 3M 500g for 100L of soil). Plants were watered every two days by an automatic watering system.

### Experimental design

Pots were haphazardly assigned to one of four experimental treatments: control, herbivory only jasmonate only, herbivory and jasmonate spray. For plants under the control treatment (C), leaves were sprayed with a sham solution containing only water and polysorbate until all leaves were wet (see Supplementary Materials for detailed solution formulae). The herbivory treatment (H) consisted of cutting off half of every second leaf on the plant with scissors and spraying plants with a sham solution until all leaves were wet (defoliation resulted in a 25% reduction of total leaf area over the course of the whole plant’s lifetime). In the jasmonate treatment (JA) plants were sprayed with a solution of methyl-jasmonate and polysorbate until all leaves were wet (polysorbate 20 was used to fix the methyl-jasmonate on the sprayed leaves). Finally, the jasmonate and herbivory treatment (JAH) consisted of cutting off half of every other leaf on the plant with scissors and spraying plants with the methyl-jasmonate solution until all leaves were wet. These treatments were applied repeatedly as plants continued to grow, i.e., they represent chronic stress or manipulation. The first round of treatment was applied one week after repotting the plants (25th of November 2019) and then every two weeks over the next 12 weeks (the last treatment was applied on the 2^nd^ of February 2020). On the first round of treatment, when most plants had fewer than six leaves each, we cut off only half a leaf (~10% of the leaf area removed) for plants under the herbivory treatments to avoid seedlings death.

Because jasmonate is highly volatile and can be perceived through stomata by neighbouring plants (Heil and Karban, 2010, Heil, 2009), plants from a given treatment were enclosed within two-metre curtains of transparent plastic to prevent ‘eavesdropping’ cross-contamination. The eight identical enclosures consisted of an aluminium squared frame (130 cm x 130 cm) to which we attached four wooden poles (two meters long). The wooden structures were wrapped with two metre-tall plastic curtains, which were fixed on three sides. The fourth side was held in place with pins acting as a door that allowed us to enter the enclosure for plant manipulations and measurements. To reduce the risk of contamination, enclosures remained closed at all times, and were opened only to allow access for measurements or manipulations. To avoid confounding effects of enclosure and treatment, we had a blocked design with two enclosures per treatment and 63 individuals per block.

### Sampling

Plant sampling consisted of cutting all above-ground plant material of 34 plants per enclosure (*N* = 272) and recording total height. Plants were then cut in half, lengthwise, creating two distinct segments: top and bottom. The top segment was carefully examined and we counted the number of fruits (immature and mature) and harvested all male flowers using tweezers. Male flowers were stored in paper envelopes, dried and weighed. After phenotyping, plant segments were dried and weighed to obtain plant dry biomass (top + bottom). To estimate seed production, the seeds were isolated from the dried plant materials, stored in paper envelopes and weighed. All materials were dried in an oven at 50°C for at least 14 days and weighed using a digital scale.

To increase the sampling efficiency, we first conducted a pilot subsampling study to test whether flower production on the top half of the plant was correlated with the total flower production of the whole plant. A strong correlation would allow considering the top plant segments as a good estimate (hereafter, subsample) of the whole plant. Plant subsampling consisted of cutting all above-ground plant material of five plants per enclosure (*N* = 40) and recording total height. The plants were cut in half, lengthwise, creating two distinct segments: top and bottom, both of which were phenotyped following the procedure detailed above. The accuracy of the top segment as a valid subsample was tested using a correlation between reproductive structures on the top segment, *versus* reproductive structures on the whole plant (top + bottom segments). We found a significant and positive correlation (*r* = 0.71; *P* = 3.94^-07^) concluding that the top sections were an accurate representation of the whole plant flower production, and proceeded with phenotyping only the top segment of the remaining 232 plants. All further statistical analyses on reproductive effort were conducted considering only plant subsample data (top segment), even for those 40 plants which were entirely phenotyped.

### Statistical analyses

All statistical analyses were conducted in *R* version 4.03 (R Core Team 2020). Mixed effects models were fitted using the ‘lme4’ package (Bates et al., 2016), residuals and model assumptions were checked using the ‘DHARMa’ package (Hartig and Hartig, 2017), and *post-hoc* Tukey comparisons were tested using the ‘multcomp’ package (Hothorn et al., 2008), and variance explained by fixed and random factors was estimated using the ‘MuMIn’ package (Barton and Barton, 2018). Model specifications, estimates and statistics are reported in Table 1. We included days-post-treatment (DPT) in our statistical models to account for the effects that the period elapsed between the last treatment application and the plant sampling date may have on our response variables. For logistical reasons, our sampling was spread over 14 days by a team of six assistants. We included observer in our statistical analyses to account for possible differences among assistants.

**Table 1.**
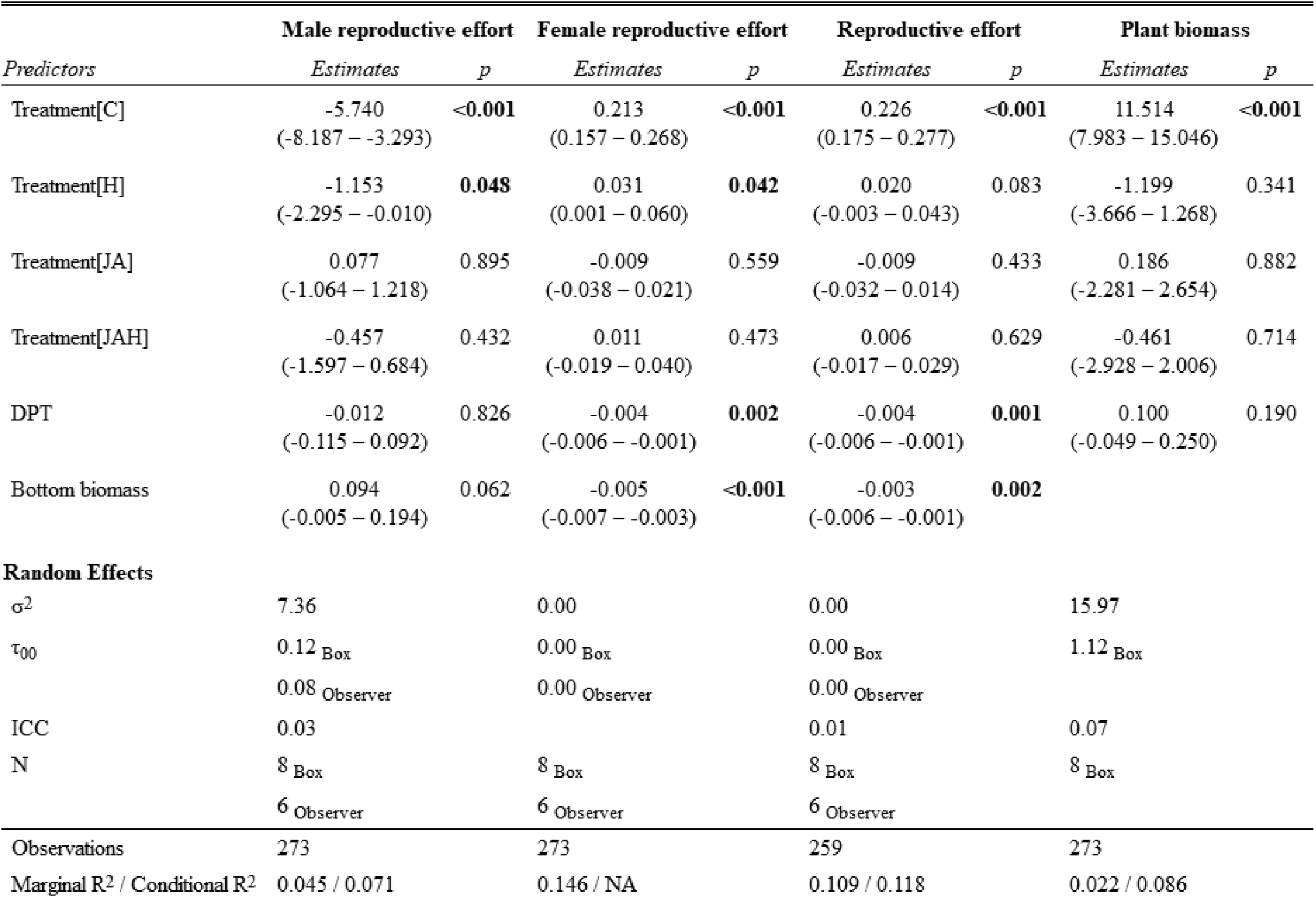
Model outputs on the effects of herbivory on male reproductive effort, female reproductive effort, total reproductive effort, and plant biomass of *Mercurialis annua*. DPT: days-post-treatment. C: control, H: herbivory (25% chronic defoliation), JA: jasmonate application, JAH: 25% chronic tissue loss and jasmonate application. Numbers in brackets show the upper and lower 95% CI of mean model estimates. Numbers in bold indicate significant differences (*P* < 0.05).

| *Predictors* | Male reproductive effort |  | Female reproductive effort |  | Reproductive effort |  | Plant biomass |  |
| --- | --- | --- | --- | --- | --- | --- | --- | --- |
|  | *Estimates* | *p* | *Estimates* | *p* | *Estimates* | *p* | *Estimates* | *p* |
| Treatment[C] | -5.740<br>(-8.187 – -3.293) | **<0.001** | 0.213<br>(0.157 – 0.268) | **<0.001** | 0.226<br>(0.175 – 0.277) | **<0.001** | 11.514<br>(7.983 – 15.046) | **<0.001** |
| Treatment[H] | -1.153<br>(-2.295 – -0.010) | **0.048** | 0.031<br>(0.001 – 0.060) | **0.042** | 0.020<br>(-0.003 – 0.043) | 0.083 | -1.199<br>(-3.666 – 1.268) | 0.341 |
| Treatment[JA] | 0.077<br>(-1.064 – 1.218) | 0.895 | -0.009<br>(-0.038 – 0.021) | 0.559 | -0.009<br>(-0.032 – 0.014) | 0.433 | 0.186<br>(-2.281 – 2.654) | 0.882 |
| Treatment[JAH] | -0.457<br>(-1.597 – 0.684) | 0.432 | 0.011<br>(-0.019 – 0.040) | 0.473 | 0.006<br>(-0.017 – 0.029) | 0.629 | -0.461<br>(-2.928 – 2.006) | 0.714 |
| DPT | -0.012<br>(-0.115 – 0.092) | 0.826 | -0.004<br>(-0.006 – -0.001) | **0.002** | -0.004<br>(-0.006 – -0.001) | **0.001** | 0.100<br>(-0.049 – 0.250) | 0.190 |
| Bottom biomass | 0.094<br>(-0.005 – 0.194) | 0.062 | -0.005<br>(-0.007 – -0.003) | **<0.001** | -0.003<br>(-0.006 – -0.001) | **0.002** |  |  |
| **Random Effects** |  |  |  |  |  |  |  |  |
| $\sigma^2$ | 7.36 | | 0.00 | | 0.00 | | 15.97 | |
| $\tau_{00}$ | 0.12 Box | | 0.00 Box | | 0.00 Box | | 1.12 Box | |
|  | 0.08 Observer |  | 0.00 Observer |  | 0.00 Observer |  |  |  |
| ICC | 0.03 |  |  |  | 0.01 |  | 0.07 |  |
| N | 8 Box |  | 8 Box |  | 8 Box |  | 8 Box |  |
|  | 6 Observer |  | 6 Observer |  | 6 Observer |  |  |  |
| Observations | 273 |  | 273 |  | 259 |  | 273 |  |
| Marginal R<sup>2</sup> / Conditional R<sup>2</sup> | 0.045 / 0.071 |  | 0.146 / NA |  | 0.109 / 0.118 |  | 0.022 / 0.086 |  |

To test the effects of herbivory on the male (MRE) and female (FRE) reproductive effort, defined as the proportion of biomass allocated to each sexual function per gram of biomass of the top subsample, we used Gaussian mixed models. To deal with zeroes and meet normality assumptions, we transformed MRE data by adding a value ten times smaller than the smallest non-zero value within its range, and applied the following formula: log(MRE) = *log*(MRE + 1^-06^). Both of these models included as fixed effects treatment, DPT, and the biomass of the bottom plant section to account for the fact that larger plants invest more in reproduction; exclusion box and observer were included as random effects.

We tested the effects that herbivory may have had on plants’ investment in sexual reproduction using a Gaussian mixed model. Reproductive effort (RE), defined as the proportion of above-ground plant biomass allocated to sexual reproduction (including male and female flowers and fruits), was fitted as the response variable. Treatment, DPT, and the biomass of the bottom plant section were fitted as fixed effects; exclusion box and observer were included as random effects. We tested whether plants responded to herbivory through compensatory growth using a Gaussian mixed model, fitting total above-ground plant biomass as the response variable, treatment and DPT as fixed effects, and the exclusion box as a random effect (for this test, we did not include observer as a random effect because all samples were weighed by one person).

## Results

Overall investment in the male function represented less than 3% of total aboveground biomass in hermaphrodites, four times less than the proportion of plant biomass invested in the female function (Fig. 1 A-B). Tissue loss significantly halved male reproductive effort in hermaphrodites compared to control plants (C: 2.47 ± 0.47; H: 1.36 ± 0.25; mean ± SE; Fig. 1A), but jasmonate application restored male reproductive investment, as shown by the lack of significant differences between JAH and control plants (Table 1, Fig. 1A). Interestingly, jasmonate alone did not enhance investment towards the male function, as shown by the non-significant differences between control plants and those only sprayed with jasmonate (Table 1, Fig. 1A). The time elapsed between the application of the last treatment and sampling date (DPT) did not significantly affect male reproductive effort (Table 1).

**Figure 1.**
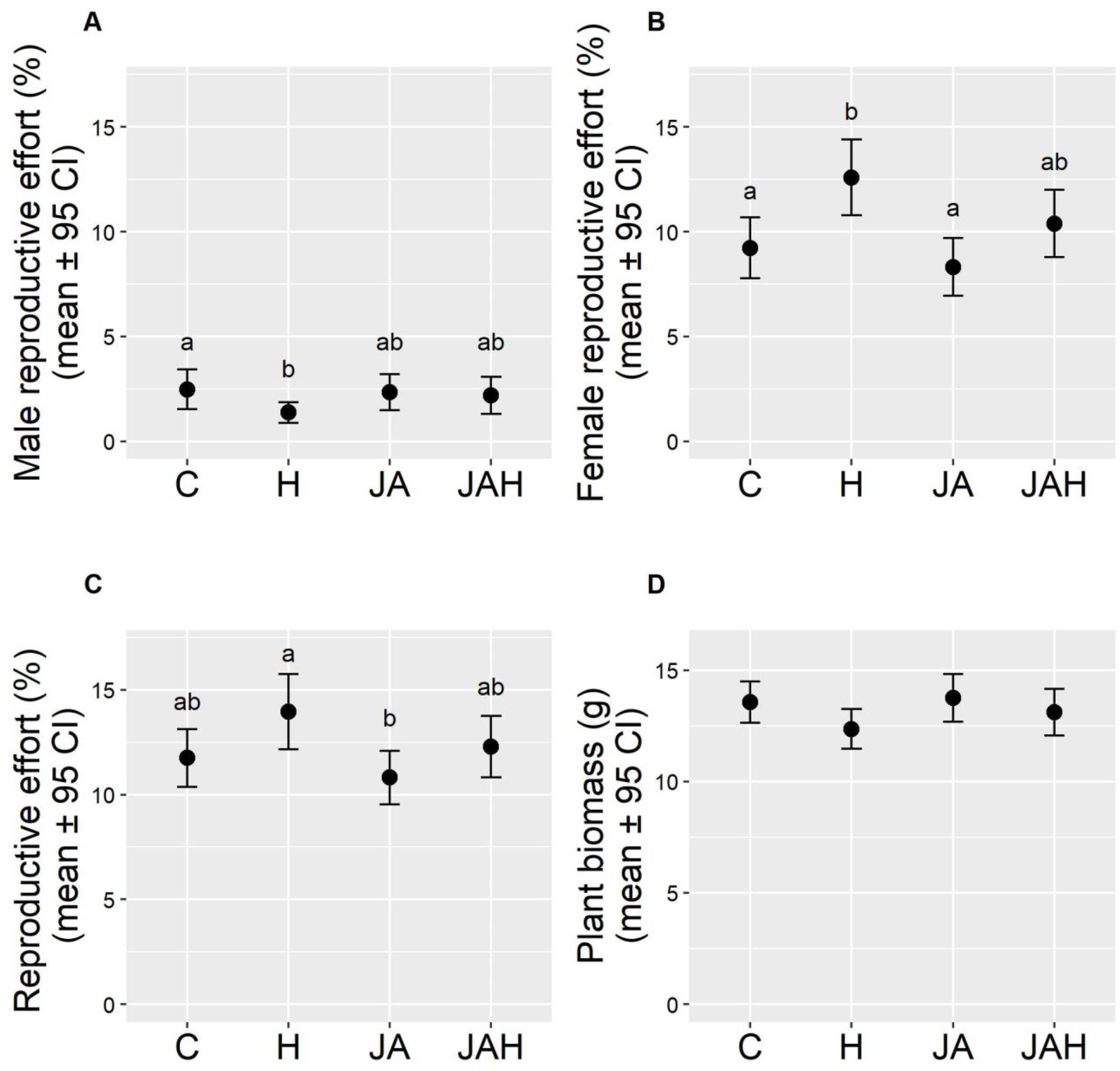
Direct (tissue loss) and indirect (jasmonate-mediated) effects of herbivory on (A) male reproductive effort (male inflorescence biomass / vegetative biomass), (B) female reproductive function (seed biomass / vegetative biomass), (C) total reproductive effort (reproductive biomass / vegetative biomass), (D) total aboveground plant biomass. Note that variables in A-C were calculated on the basis of measurements of the apical subsample for each individual sampled. Treatments are abbreviated as follows, C: Control, H: herbivory treatment with a chronic tissue loss of 25% of foliar area, JA: jasmonate application; JAH: tissue loss and jasmonate application treatment. Different letters indicate significant differences between treatment levels (*P* < 0.05). Total *N* = 273 plants.

Tissue loss significantly increased female reproductive effort in hermaphrodites by a quarter compared to control plants or those only sprayed by jasmonate (C: 9.22 ± 0727; H: 12.58 ± 0.90; JA: 8.30 ± 0.69; mean ± SE; Fig. 1B), but, again, jasmonate application in combination with tissue loss restored reproductive investment to levels similar to those of control plants, as shown by the lack of significant differences between JAH and control plants (Table 1, Fig. 1B). The time elapsed between the application of the last treatment and sampling date (DPT) had a negative effect on female reproductive effort (Table 1). In all models, random effects explained very little of the variance of the response variables, indicating that the enclosure box and the observer had a negligible effect on our estimates (Table 1).

The herbivory treatments had no significant effect on plant biomass. Despite suffering 25% defoliation, plants under the herbivory treatment were on average only 8.85% lighter than control plants (C: 13.56 ± 0.46; H: 12.36 ± 0.44; mean ± SE; Fig. 1D), a difference that was statistically non-significant (Table 1, Fig 1D). The time elapsed between the application of the last treatment and sampling date (DPT) did not significantly affect plant biomass (Table 1). On average, approximately 25% of reproductive biomass was allocated to male flower production, whilst the remaining 75% was allocated towards the female function. Yet, neither tissue loss nor jasmonate application had a significant effect on the proportion of biomass allocated towards sexual reproduction (reproductive effort; Table 1, Fig. 1C). Plants invested on average ~10% of their total aboveground biomass towards reproduction, even under chronic 25% defoliation. Similarly, jasmonate application had no significant effect on reproductive effort (Fig. 1C). The time elapsed between the application of the last treatment and sampling date (DPT) had a negative effect on reproductive effort (Table 1).

## Discussion

Even though chronic 25% defoliation had no significant effect on plant biomass, tissue loss significantly increased the proportion of biomass allocated towards reproduction. This increase in reproductive effort was invested towards the female function, at the expense of investment in the male function. However, exogenous jasmonate application effectively restored investment towards the male function in damaged plants.

### Herbivory-induced sex allocation was altered by selection in the absence of herbivory

Our experiment revealed that tissue loss due to chronic simulated herbivory in *M. annua* caused females with enhanced leaky sex expression in response to recent selection (Cossard et al., 2021) to shift their sex allocation away from male flower production. This response was thus opposed to that of their less leaky recent ancestors, for which tissue damage *increased* male flower production (Villamil et al., 2021). In other words, the very direction of the reaction norm of male flower production by females in response to tissue damage in natural populations of dioecious *M. annua* has been altered by strong selection on male flower production.

We do not know whether the reaction norm of male-flower production by females in the ancestral dioecious populations is adaptive, but it might be so (Villamil et al., 2021). However, we can safely say that the change in reaction norm that we have observed cannot be adaptive, for two reasons. First, females evolved enhanced male production in the selection experiment in the complete absence of sustained tissue damage (and, more importantly in the absence of variation in tissue damage), so they were not exposed to selection that might have favoured a different reaction norm. Second, the mean male flower production in the females of our experiment was much higher than is ever found in natural populations, such that, again, reaction norms around this new mean are not pertinent to the optimising effects of natural selection in the ancestral population. The reaction norms we have measured are therefore, almost by definition, non-adaptive, and should properly be seen as a pleiotropic effect of selection on reproductive allocation.

Pleiotropic effects of natural selection have been found in selection experiments in a number of contexts (Hill and Caballero, 1992) from theoretical models (Otto, 2004) to bacteria (Lenski, 1988a, Lenski, 1988b, Jasmin and Zeyl, 2013), insects (Harshman and Hoffmann, 2000), or plants. A classic example of this are Lenski’s *Escherichia coli* lines, which evolved resistance to a virus along with maladaptive mutations on various metabolic pathways and which eventually resulted in reduced fitness levels, similar to those in non-resistant populations (Lenski, 1988a, Lenski, 1988b). A more recent example in *Drosphila* shows that evolved resistance against an intestinal pathogen evolved along with deleterious effects on male fitness through reduced male mating success and reduced sperm competitive abilities (Kawecki, 2020). In plants, maladaptive pleiotropic effects in response to an experimental selection pressure have been widely observed in the context of evolution of herbicide resistance (Bergelson et al., 1996, Debban et al., 2015), but also in that of the evolution of flower colour (Coberly and Rausher, 2008), induced defences (Siemens and Mitchell-Olds, 1998), flowering phenology (Kover et al., 2009), or increased plant size leading to decreased defensive strategies and reduced floral signals (Zu and Schiestl, 2017). Our study provides a further revealing illustration of the complexity of multi-trait responses to selection, and, notably, of the fact that quantitative genetic G-matrices include not only simple traits but also environmentally dependent reaction norms.

### Tissue loss caused plants to reduce male sex allocation, but jasmonate signalling restored it

Our experiment tested both the direct (tissue loss) and indirect (jasmonate signalling) effects of herbivory on sex allocation in the evolved population of *M. annua*. Interestingly, we found that plants reacted to both of these components differently. Male reproductive effort was nearly halved in defoliated plants, and jasmonate restored male investment after tissue loss, showing that signalling related to anti-herbivore defences might also affect sexual expression. We expected the tissue loss and jasmonate application treatments to have additive effects on male investment, with the exogenous (jasmonate application) signal effectively amplifying the endogenous direct effects of tissue loss. However, jasmonate signalling caused plants to increase male net investment only if they had suffered tissue loss. To some extent, this result contrasts with previous findings showing that jasmonate can induce plant responses even in the absence of tissue loss (Hernandez-Cumplido et al., 2016, Heil et al., 2001b, Heil, 2015, Kost and Heil, 2008, Radhika et al., 2008, Heil et al., 2001a, Escalante-Pérez and Heil, 2012, Kessler and Heil, 2011). If jasmonate on its own enhanced male allocation, whilst tissue loss reduced it, we expected plants that only received additional jasmonate but suffered no tissue loss (JA) to have greater male investment than those under the tissue loss and jasmonate treatment (JAH). However, we found no differences between plants in J and JAH treatments in male investment or any of the response variables measured. Our results thus show that an increased jasmonate signal perceived exogenously can restore male sex allocation only in damaged plants. As explained above, it is difficult to infer any functional or adaptive explanation for this result, but it is clear that different effects of damage interact and are not simply additive.

The exogenous application of jasmonate that restored male sex allocation to non-herbivory levels would be a favourably selected response in the recent evolutionary past of the *M. annua* population (in which males had been removed), because the male-flower production provides exceptionally high fitness through siring (Cossard and Pannell, 2021, Cossard et al., 2021). The ability of jasmonate to restore male sex allocation when plants are damaged may thus also be a pleiotropic effect of selection, perhaps as a result of a physiological constraint associated with hormonal cross-talk. Clearly, with time and appropriate conditions, we should expect natural selection to act upon the reaction norms we have observed, gradually modifying the non-adaptive response to jasmonate in restoring male allocation and converting it to a response that could legitimately be seen as adaptive. For instance, this could occur if plants evolved increased sensitivity to jasmonate as a male-function restorer so that the smaller amounts of jasmonate released exclusively by damage would suffice to restore male sex allocation in defoliated plants. Such an evolved response would allow damaged females with enhanced leakiness (H) to maximise their fitness, regardless of the herbivory levels experienced by their neighbours, i.e., the additional jasmonate levels. Future research in other species with a longer evolutionary history under monoecy and under suitable variation in herbivory levels would be valuable to determine whether jasmonate is also capable of restoring tissue loss-induced shifts in sex expression and what its adaptive or deleterious consequences might be.

Our findings suggest that jasmonate signalling may connect defensive and reproductive strategies in *M. annua*. More specifically, they indicate that herbivory might affect sex expression through its direct effects on tissue loss, but not through its indirect influence on hormone signalling, as perceived from neighbouring plants (by eavesdropping), and that jasmonate perceived in synergy with damage might restore the effects of tissue loss. Jasmonate has been found to play a role in plant responses to herbivory (Ballaré, 2011, Thaler et al., 2001, Heil and Bueno, 2007b, Heil and Karban, 2010) and in regulating sex expression, flower development and sexual differentiation in a number of other plants species (Wasternack et al., 2013, Yuan and Zhang, 2015, Cai et al., 2014, Acosta et al., 2009, Yan et al., 2012), and our results now confirm that it can act upon both functions simultaneously in *M. annua*; to our knowledge, this is the first account of the influence of jasmonate on the sexual expression of a species in the Euphorbiaceae family. Even though *M. annua* has chromosomal sex determination (Durand and Durand, 1991, Russell and Pannell, 2015), its sexual expression can be influenced and even switched via the exogenous application of phytohormones such as cytokinins and auxins (Durand and Durand, 1991). The pleiotropic effects of herbivory on sex expression observed in *M. annua* could well be due to hormonal ‘cross-talk’ of jasmonate with sex-determining hormones cytokinins and auxins, which has been well documented to occur in other species (Robert-Seilaniantz et al., 2011, Naseem et al., 2015).

### Tissue loss increased female sex allocation

Chronic defoliation elicited a change in male-flower production in *M. annua* females, and this change was associated with a corresponding response in fruit production, through a sex-allocation trade-off. This pattern strongly contrasts with the available evidence on monoecious species in which defoliation had the opposite effect, i.e., reducing female, but not male allocation (Snyder, 1993, Quesada et al., 1995). Furthermore, our findings also contrast with previous studies in hermaphroditic species in which defoliation decreased male sex allocation, whilst female sex allocation remained unaffected (Lehtilä and Strauss, 1997, Allison, 1990, Frazee and Marquis, 1994). In this regard, our experimental population of monoecious *M. annua* plants seems to have a herbivory-induced sex allocation response different from those previously reported in the literature for hermaphroditic or monoecious species. As noted above, this unusual pattern may be attributable to the fact that it has not yet been moulded by natural selection.

### No measurable negative impacts on performance and increased fitness: tolerance?

Contrary to our expectations, tissue loss increased the proportion of biomass invested towards sexual reproduction. Despite a chronic 25% defoliation through the plants’ lifetime, we observed no trade-offs between growth and reproduction. Rather, we found that defoliated plants significantly increased their reproductive effort by 2% compared to control plants. Plants are known to respond to damage by herbivores in three main ways: by showing ‘resistance’ (i.e., by investing in physical or chemical means to deter consumption); by ‘escaping’ (i.e., evading attack through phenological mismatching); and/or through ‘tolerance’ (i.e., compensating tissue loss with new growth) (Boege et al., 2007, Stamp, 2003, Fornoni, 2011). Our experimental plants might have coped with damage through compensatory growth, given that we observed no measurable negative impacts of defoliation on performance (biomass) (Fig. 1D). We conducted a power analysis using the *R* package ‘simr’ (Green and MacLeod, 2016) and found that, given our sample size (*N* = 272), our experiment had sufficient power (80% power) to detect effect sizes ≥ 25% on plant biomass, i.e., only much greater than the effect sizes from our experiment (−8.5% and +1.3%). Thus, while it is possible that plants compensated for and tolerated the lost tissue, our experiment lacked power to rule out substantial, non-compensatory, negative effects of herbivory.

### Concluding remarks

Taken together, our results indicate that tissue loss and jasmonate signalling both had cascading effects on sex allocation in *M. annua*, and that jasmonate can be involved in both sex expression and defence pathways simultaneously. Our results thus suggest that defence-related traits – such as those induced by jasmonate signalling – may link the evolution of reproductive and defensive traits, and that jasmonate may thus play an important role in conditional sex allocation in some plants. Conditional sex allocation provides an ideal theoretical framework to address knowledge gaps on the effects of plant defences on trade-offs between male and female plant fitness and sex allocation, advancing our understanding on the role antagonists may have as selective agents shaping sex allocation trade-offs. Appropriate experimental evolution studies would be a valuable tool to test the potential of selection to optimise this response.

## Acknowledgements

We thank Aline Revel for assistance with growing plants, and Juan Fornoni and Kai-Hsiu Chen for comments that improved earlier versions of this manuscript. Funders: Swiss National Science Foundation (grant 31003A_163384 to JRP) and the University of Lausanne.

## SUPPLEMENTARY MATERIAL

### Appendix 1: Protocol for preparing sham and methyl-jasmonate solutions

Methyl-jasmonate solution:

1. 1 mL of Polysorbate 20 (tween20) gauged to 250 mL of distilled water
2. 0.1 mL of Methyl-jasmonate gauged to 250 mL of distilled water
3. Mix both solution, gauge to 1000 mL and transfer to a jar with an airtight lid.
4. Vortex the mix 45 sec (closed jar to avoid volatiles escaping) and keep the solution at 4°C.

Sham solution used for the control treatment.

1. 1 mL of Polysorbate 20 (tween20) gauged to 1000 mL of distilled water.
2. Transfer to a jar with an airtight lid.
3. Vortex the mix 45 sec (closed jar) and keep the solution at 4°C.

## Notes

### Competing Interest Statement

The authors have declared no competing interest.

## References

Acosta IF, Laparra H, Romero SP, Schmelz E, Hamberg M, Mottinger JP, Moreno MA, Dellaporta SL. 2009. tasselseed1 is a lipoxygenase affecting jasmonic acid signaling in sex determination of maize. Science, 323: 262–265.

Allison TD. 1990. The influence of deer browsing on the reproductive biology of Canada yew (Taxus canadensis marsh.). Oecologia, 83: 523–529.

Anastasaki E, Balayannis G, Papanikolaou NE, Michaelakis AN, Milonas PG. 2015. Oviposition induced volatiles in tomato plants. Phytochemistry Letters, 13: 262–266.

Ballaré CL. 2011. Jasmonate-induced defenses: a tale of intelligence, collaborators and rascals. Trends in Plant Science, 16: 249–257.

Barrett SC. 2002. The evolution of plant sexual diversity. Nature Reviews Genetics, 3: 274–284.

Barton K, Barton M. 2018. Package ‘MuMIn’. R package version 3.1.

Bates D, Maechler M, Bolker B, Walker S, Christensen RHB, Singmann H, Dai B, Grothendieck G, Green P, Bolker MB. 2016. Package ‘lme4’. R Package Version 1.1–10.

Bergelson J, Purrington CB, Palm CJ, López-Gutiérrez J-c. 1996. Costs of resistance: a test using transgenic <i>Arabidopsis thaliana</i>. Proceedings of the Royal Society of London. Series B: Biological Sciences, 263: 1659–1663.

Bierzychudek P. 1984a. Assessing “optimal” life histories in a fluctuating environment: the evolution of sex-changing by jack-in-the-pulpit. The American Naturalist, 123: 829–840.

Bierzychudek P. 1984b. Determinants of gender in Jack-in-the-pulpit: the influence of plant size and reproductive history. Oecologia, 65: 14–18.

Boege K, Dirzo R, Siemens D, Brown P. 2007. Ontogenetic switches from plant resistance to tolerance: minimizing costs with age? Ecology Letters, 10: 177–187.

Browse J. 2009. The power of mutants for investigating jasmonate biosynthesis and signaling. Phytochemistry, 70: 1539–1546.

Cai Q, Yuan Z, Chen M, Yin C, Luo Z, Zhao X, Liang W, Hu J, Zhang D. 2014. Jasmonic acid regulates spikelet development in rice. Nature Communications, 5: 1–13.

Campbell S, Kessler A. 2013. Plant mating system transitions drive the macroevolution of defense strategies. Proceedings of the National Academy of Sciences of the United States of America, 110: 3973–3978.

Campbell SA. 2015. Ecological mechanisms for the coevolution of mating systems and defence. New Phytologist, 205: 1047–1053.

Carr DE, Eubanks MD. 2014. Interactions between insect herbivores and plant mating systems. Annual Review of Entomology, 59: 185–203.

Charnov EL. 1982. The Theory of Sex Allocation: Princeton University Press.

Coberly LC, Rausher MD. 2008. Pleiotropic effects of an allele producing white flowers in *Ipomoea purpurea*. Evolution, 62: 1076–1085.

Cossard GG, Gerchen JF, Li X, Cuenot Y, Pannell JR. 2021. The rapid dissolution of dioecy by experimental evolution. Current Biology, 31: 1277–1283.

Cossard GG, Pannell JR. 2019. A functional decomposition of sex inconstancy in the dioecious, colonizing plant Mercurialis annua. American Journal of Botany, 106: 722–732.

Cossard GG, Pannell JR. 2021. Enhanced leaky sex expression in response to pollen limitation in the dioecious plant Mercurialis annua. Journal of Evolutionary Biology, 34: 416–422.

de Jong T, Klinkhamer P. 1989. Size-dependency of sex-allocation in hermaphroditic, monocarpic plants. Functional Ecology, 3: 201–206.

de Jong T, Klinkhamer P. 2005. Evolutionary ecology of plant reproductive strategies: Cambridge University Press.

de Jong TJ, Waser NM, Price MV, Ring RM. 1992. Plant size, geitonogamy and seed set in Ipomopsis aggregata. Oecologia, 89: 310–315.

Debban CL, Okum S, Pieper KE, Wilson A, Baucom RS. 2015. An examination of fitness costs of glyphosate resistance in the common morning glory, Ipomoea purpurea. Ecology and Evolution, 5: 5284–5294.

Durand B, Durand R. 1991. Sex determination and reproductive organ differentiation in Mercurialis. Plant Science, 80: 49–65.

Ehlers BK, Bataillon T. 2007. ‘Inconstant males’ and the maintenance of labile sex expression in subdioecious plants. New Phytologist, 174: 194–211.

Escalante-Pérez M, Heil M. 2012. Nectar secretion: its ecological context and physiological regulation. In: Vivanco J. BF, ed. Secretions and exudates in biological systems. Berlin, Heidelberg: Springer-Verlag.

Farmer EE, Gao YQ, Lenzoni G, Wolfender JL, Wu Q. 2020. Wound-and mechanostimulated electrical signals control hormone responses. New Phytologist, 227: 1037–1050.

Fornoni J. 2011. Ecological and evolutionary implications of plant tolerance to herbivory. Functional Ecology, 25: 399–407.

Frazee JE, Marquis RJ. 1994. Environmental contribution to floral trait variation in Chamaecrista fasciculata (Fabaceae: Caesalpinoideae). American Journal of Botany, 81: 206–215.

Freeman D, Harper K, Charnov EL. 1980. Sex change in plants: old and new observations and new hypotheses. Oecologia, 47: 222–232.

Garcia LC, Eubanks MD. 2019. Overcompensation for insect herbivory: a review and meta-analysis of the evidence. Ecology, 100: e02585.

Ghiselin MT. 1969. The evolution of hermaphroditism among animals. The Quarterly Review of Biology, 44: 189–208.

Green P, MacLeod CJ. 2016. SIMR: an R package for power analysis of generalized linear mixed models by simulation. Methods in Ecology and Evolution, 7: 493–498.

Hambäck PA. 2001. Direct and indirect effects of herbivory: feeding by spittlebugs affects pollinator visitation rates and seedset of *Rudbeckia hirta*. Ecoscience, 8: 45–50.

Han G-Z. 2016. Evolution of jasmonate biosynthesis and signaling mechanisms. Journal of Experimental Botany, 68: 1323–1331.

Harshman LG, Hoffmann AA. 2000. Laboratory selection experiments using Drosophila: what do they really tell us? Trends in Ecology & Evolution, 15: 32–36.

Hartig F, Hartig MF. 2017. Package ‘DHARMa’: residual diagnostics for hierarchical (multi-level/mixed) regression models. R package..

Heil M. 2009. Airborne induction and priming of defenses. Plant-Environment Interactions: Springer.

Heil M. 2015. Extrafloral nectar at the plant-insect interface: a spotlight on chemical ecology, phenotypic plasticity, and food webs. Annual Review of Entomology, 60: 213–232.

Heil M, Brigitte F, Ulrich M, Linsenmair KE. 2001a. On benefits of indirect defence: short- and long-term studies of antiherbivore protection via mutualistic ants. Oecologia, 126.

Heil M, Bueno JCS. 2007a. Herbivore-induced volatiles as rapid signals in systemic plant responses: how to quickly move the information? Plant signaling & behaviour, 2: 191–193.

Heil M, Bueno JCS. 2007b. Within-plant signaling by volatiles leads to induction and priming of an indirect plant defense in nature. Proceedings of the National Academy of Sciences, 104: 5467–5472.

Heil M, Karban R. 2010. Explaining evolution of plant communication by airborne signals. Trends in ecology & evolution, 25: 137–144.

Heil M, Koch T, Hilpert A, Fiala B, Boland W, Linsenmair KE. 2001b. Extrafloral nectar production of the ant-associated plant, Macaranga tanarius, is an induced, indirect, defensive response elicited by jasmonic acid. Proceedings of the National Academy of Sciences, 98: 1083–1088.

Hendrix SD, Trapp EJ. 1981. Plant-herbivore interactions: insect induced changes in host plant sex expression and fecundity. Oecologia, 49: 119–122.

Herms DA, Mattson WJ. 1992. The dilemma of plants: to grow or defend. Quarterly review of biology: 283–335.

Hernandez-Cumplido J, Forter B, Moreira X, Heil M, Benrey B. 2016. Induced Floral and Extrafloral Nectar Production Affect Ant-pollinator Interactions and Plant Fitness. Biotropica.

Hilker M, Meiners T. 2002. Induction of plant responses to oviposition and feeding by herbivorous arthropods: a comparison. Proceedings of the 11th International Symposium on Insect-Plant Relationships: Springer.

Hill WG, Caballero A. 1992. Artificial selection experiments. Annual Review of Ecology and Systematics, 23: 287–310.

Hirata R, Wasaka N, Fujii A, Kato T, Sato H. 2019. Differences in flowering phenology, architecture, sexual expression and resource allocation between a heavily haired and a lightly haired nettle population: relationships with sika deer. Plant Ecology, 220: 255–266.

Hothorn T, Bretz F, Westfall P, Heiberger R. 2008. Multcomp: simultaneous inference for general linear hypotheses. R Package Version 1.0–3.

Ivey CT, Carr DE. 2005. Effects of herbivory and inbreeding on the pollinators and mating system of Mimulus guttatus (Phrymaceae). American Journal of Botany, 92: 1641–1649.

Jasmin J-N, Zeyl C. 2013. Evolution of pleiotropic costs in experimental populations. Journal of Evolutionary Biology, 26: 1363–1369.

Johnson MT, Campbell SA, Barrett SC. 2015. Evolutionary interactions between plant reproduction and defense against herbivores. Annual Review of Ecology, Evolution, and Systematics, 46: 191–213.

Karban R. 2008. Plant behaviour and communication. Ecology letters, 11: 727–739.

Karban R, Ishizaki S, Shiojiri K. 2012. Long-term demographic consequences of eavesdropping for sagebrush. Journal of Ecology, 100: 932–938.

Karban R, Strauss SY. 1993. Effects of herbivores on growth and reproduction of their perennial host, *Erigeron glaucus*. Ecology, 74: 39–46.

Kawecki TJ. 2020. Sexual selection reveals a cost of pathogen resistance undetected in life-history assays. Evolution, 74: 338–348.

Kessler A, Halitschke R, Poveda K. 2011. Herbivory-mediated pollinator limitation: Negative impacts of induced volatiles on plant-pollinator interactions. Ecology, 92: 1769–1780.

Kessler A, Heil M. 2011. The multiple faces of indirect defences and their agents of natural selection. Functional Ecology, 25.

Klinkhamer PG, De Jong TJ, Metz H. 1997. Sex and size in cosexual plants. Trends in Ecology & Evolution, 12: 260–265.

Kost C, Heil M. 2008. The defensive role of volatile emission and extrafloral nectar secretion for lima bean in nature. Journal of Chemical Ecology, 34: 1–13.

Kover PX, Rowntree JK, Scarcelli N, Savriama Y, Eldridge T, Schaal BA. 2009. Pleiotropic effects of environment-specific adaptation in Arabidopsis thaliana. New Phytologist, 183: 816–825.

Krupnick G, Avila G, Brown K, Stephenson A. 2000. Effects of herbivory on internal ethylene production and sex expression in Cucurbita texana. Functional Ecology: 215–225.

Krupnick GA, Weis AE. 1998. Floral herbivore effect on the sex expression of an andromonoecious plant, Isomeris arborea (Capparaceae). Plant Ecology, 134: 151–162.

Lehtilä K, Strauss SY. 1997. Leaf damage by herbivores affects attractiveness to pollinators in wild radish, *Raphanus raphanistrum*. Oecologia, 111: 396–403.

Lehtilä K, Strauss SY. 1999. Effects of foliar herbivory on male and female reproductive traits of wild radish, *Raphanus raphanistrum*. Ecology, 80: 116–124.

Lenski RE. 1988a. Experimental studies of pleiotropy and epistais in Escherichia coli I. Variation in competitive fitness among mutants resistant to virus T4.. Evolution, 42: 425–432.

Lenski RE. 1988b. Experimental studies of pleiotropy and epistasis in *Escherichia coli.* II. Compensation for maladaptive effects associated with resistance to virus T4. Evolution, 42: 433–440.

Li L, Zhao Y, McCaig BC, Wingerd BA, Wang J, Whalon ME, Pichersky E, Howe GA. 2004. The tomato homolog of CORONATINE-INSENSITIVE1 is required for the maternal control of seed maturation, jasmonate-signaled defense responses, and glandular trichome development. The Plant Cell, 16: 126–143.

López S, Domínguez C. 2003. Sex choice in plants: facultative adjustment of the sex ratio in the perennial herb Begonia gracilis. Journal of Evolutionary Biology, 16: 1177–1185.

Lucas-Barbosa D. 2016. Integrating studies on plant-pollinator and plant-herbivore interactions. Trends in plant science, 21: 125–133.

Mothershead K, Marquis RJ. 2000. Fitness impacts of herbivory through indirect effects on plant-pollinator interactions in Oenothera macrocarpa. Ecology, 81: 30–40.

Mumm R, Schrank K, Wegener R, Schulz S, Hilker M. 2003. Chemical analysis of volatiles emitted by Pinus sylvestris after induction by insect oviposition. Journal of chemical ecology, 29: 1235–1252.

Naseem M, Kaltdorf M, Dandekar T. 2015. The nexus between growth and defence signalling: auxin and cytokinin modulate plant immune response pathways. Journal of Experimental Botany, 66: 4885–4896.

Obbard DJ, Harris SA, Buggs RJ, Pannell JR. 2006. Hybridization, polyploidy, and the evolution of sexual systems in Mercurialis (Euphorbiaceae). Evolution, 60: 1801–1815.

Otto SP. 2004. Two steps forward, one step back: the pleiotropic effects of favoured alleles. Proceedings of the Royal Society of London. Series B: Biological Sciences, 271: 705–714.

Pannell J. 1997a. Mixed genetic and environmental sex determination in an androdioecious population of Mercurialis annua. Heredity, 78: 50–56.

Pannell J. 1997b. Variation in sex ratios and sex allocation in androdioecious Mercurialis annua. Journal of Ecology: 57–69.

Pannell JR, Dorken ME, Pujol B, Berjano R. 2008. Gender variation and transitions between sexual systems in Mercurialis annua (Euphorbiaceae). International Journal of Plant Sciences, 169: 129–139.

Policansky D. 1987. Sex choice and reproductive costs in jack-in-the-pulpit. Bioscience, 37: 476–481.

Poveda K, Steffan-Dewenter I, Scheu S, Tscharntke T. 2003. Effects of below-and above-ground herbivores on plant growth, flower visitation and seed set. Oecologia, 135: 601–605.

Quesada M, Bollman K, Stephenson AG. 1995. Leaf damage decreases pollen production and hinders pollen performance in Cucurbita texana. Ecology, 76: 437–443.

Radhika V, Kost C, Bartram S, Heil M, Boland W. 2008. Testing the optimal defence hypothesis for two indirect defences: extrafloral nectar and volatile organic compounds. Planta, 228: 449–457.

Ramos SE, Schiestl F. 2019. Rapid plant evolution driven by the interaction of pollination and herbivory. Science, 364: 193–196.

Reymond P, Farmer EE. 1998. Jasmonate and salicylate as global signals for defense gene expression. Current opinion in plant biology, 1: 404–411.

Robert-Seilaniantz A, Grant M, Jones JDG. 2011. Hormone crosstalk in plant disease and defense: more than just jasmonate-salicylate antagonism. Annual Review of Phytopathology, 49: 317–343.

Russell J, Pannell J. 2015. Sex determination in dioecious Mercurialis annua and its close diploid and polyploid relatives. Heredity, 114: 262–271.

Salvador-Recatalà V, Tjallingii WF, Farmer EE. 2014. Real-time, in vivo intracellular recordings of caterpillar-induced depolarization waves in sieve elements using aphid electrodes. New Phytologist, 203: 674–684.

Santangelo JS, Thompson KA, Johnson MT. 2019. Herbivores and plant defences affect selection on plant reproductive traits more strongly than pollinators. Journal of Evolutionary Biology, 32: 4–18.

Siemens DH, Mitchell-Olds T. 1998. Evolution of pest-induced defenses in Brassica plants: Tests of theory. Ecology, 79: 632–646.

Snyder MA. 1993. Interactions between Abert’s squirrel and ponderosa pine: the relationship between selective herbivory and host plant fitness. The American Naturalist, 141: 866–879.

Stamp N. 2003. Out of the quagmire of plant defense hypotheses. The Quarterly review of biology, 78: 23–55.

Strauss SY, Conner JK, Rush SL. 1996. Foliar herbivory affects floral characters and plant attractiveness to pollinators: implications for male and female plant fitness. American Naturalist, 147: 1098–1107.

Strauss SY, Irwin RE, Lambrix VM. 2004. Optimal defence theory and flower petal colour predict variation in the secondary chemistry of wild radish. Journal of Ecology, 92: 132–141.

Thaler JS, Stout MJ, Karban R, Duffey SS. 2001. Jasmonate-mediated induced plant resistance affects a community of herbivores. Ecological Entomology, 26: 312–324.

Thomson V, Nicotra A, Cunningham S. 2004. Herbivory differentially affects male and female reproductive traits of Cucumis sativus. Plant Biology, 6: 621–628.

Tutin T, Heywood V, Burges N, Moore D, Valentine D, Walters S, Webb D. 1968. Flora Europaea, vol. 2-5. Cambridge University Press, Cambridge.

Vaidya P, McDurmon A, Mattoon E, Keefe M, Carley L, Lee CR, Bingham R, Anderson JTJNP. 2018. Ecological causes and consequences of flower color polymorphism in a self-pollinating plant (*Boechera stricta*). New Phytologist, 218: 380–392.

Villamil N, Li X, Seddon E, Pannell J. 2021. Herbivory enhances leaky sex expression in the dioecious herb Mercurialis annua. Annals of Botany, Accepted, in press..

Wang J, Wu D, Wang Y, Xie D. 2019. Jasmonate action in plant defense against insects. Journal of Experimental Botany, 70: 3391–3400.

Wang Y, Luo A, Lyu T, Dimitrov D, Xu X, Freckleton RP, Li Y, Su X, Li Y, Liu Y. 2021. Global distribution and evolutionary transitions of angiosperm sexual systems. Ecology Letters.

Wasternack C, Forner S, Strnad M, Hause B. 2013. Jasmonates in flower and seed development. Biochimie, 95: 79–85.

West S. 2009. Sex allocation: Princeton University Press.

Yampolsky C. 1919. Inheritance of sex in Mercurialis annua. American Journal of Botany: 410–442.

Yampolsky C. 1930. Induced alteration of sex in the male plant of *Mercurialis annua*. Bulletin of the Torrey Botanical Club: 51–58.

Yan Y, Christensen S, Isakeit T, Engelberth J, Meeley R, Hayward A, Emery RN, Kolomiets MV. 2012. Disruption of OPR7 and OPR8 reveals the versatile functions of jasmonic acid in maize development and defense. The Plant Cell, 24: 1420–1436.

Yuan Z, Zhang D. 2015. Roles of jasmonate signalling in plant inflorescence and flower development. Current opinion in plant biology, 27: 44–51.

Zu P, Schiestl FP. 2017. The effects of becoming taller: direct and pleiotropic effects of artificial selection on plant height in Brassica rapa. The Plant Journal, 89: 1009–1019.

